# Using near-infrared spectroscopy to discriminate closely related species: A case study of neotropical ferns

**DOI:** 10.1101/2020.10.19.343947

**Authors:** Darlem Nikerlly Amaral Paiva, Ricardo de Oliveira Perdiz, Thaís Elias Almeida

## Abstract

Identifying plant species requires considerable knowledge and can be difficult without complete specimens. Fourier-transform near-infrared spectroscopy (FT-NIR) is an effective technique for discriminating plant species, especially angiosperms. However, its efficacy has never been tested on ferns. Here we tested the accuracy of FT-NIR at discriminating species of the genus *Microgramma*. We obtained 16 spectral readings per individual from the adaxial and abaxial surfaces of 100 specimens belonging to 13 species. The analyses included all 1557 spectral variables. We tested different datasets (adaxial+abaxial, adaxial, and abaxial) to compare the correct identification of species through the construction of discriminant models (LDA, PLS) and cross-validation techniques (leave-one-out, K-fold). All analyses recovered an overall high percentage (>90 %) of correct predictions of specimen identifications for all datasets, regardless of the model or cross-validation used. On average, there was > 95 % accuracy when using PLS-DA and both cross-validations. Our results show the high predictive power of FT-NIR at correctly discriminating fern species when using leaves of dried herbarium specimens. The technique is sensitive enough to reflect species delimitation problems and possible hybridization, and it has the potential of helping better delimit and identify fern species.

## INTRODUCTION

Defining and identifying species using qualitative morphological traits can be challenging even though species identification is fundamental to some areas of science and sustainable dynamics (Galtier 2018; Pinheiro et al. 2018). Correct identifications also contribute significantly to understanding the evolutionary history of many species and the diversity of biological groups in rich and threatened areas, such as tropical forests (Costello 2015). Considering the biological and historical diversity of polymorphisms in plants, allied with centuries of describing species using alpha taxonomy tools, the correct identification of a specimen requires experts with considerable knowledge (Ahrends et al. 2011; Lacerda and Nimmo 2010; Richard and Evans 2006).

A problem when identifying plant species is the absence of complete specimens, including both sterile and fertile material, such as flowers or fruits of seed plants (Gomes et al. 2013). Difficult to access and insufficient or unrepresentative collections of species widely distributed in highly diverse areas can also pose a problem when identifying specimens (Lacerda and Nimmo 2010). Among the traditional identification methods used for plants, keys stand out and are widely employed (Smith 2017). However, polymorphisms and the complexity of shapes, associated with homoplasies and cryptic taxa, for example, create the need for more elaborate tools aimed primarily at the identification, conservation, and elucidation of unclear relationships of plants (Durgante et al. 2013; Pinheiro et al. 2018).

In addition to the use of macromorphology, DNA barcoding is an internationally recognized tool and widely used in species identification, ecological studies, and forensic analyses (Li et al. 2015; Shokralla et al. 2014). In studies of animal groups that used this molecular approach, the technique proved to be highly efficient (e.g., Ohira et al. 2018; Porco et al. 2012; Pérez-Losada et al. 2012). Using DNA barcoding has been less successful at identifying plants compared to animals (Li et al. 2015). According to Fazekas et al. (2012), this is partially due to hybridization, polyploidy, and speciation related to reproductive systems. However, these are not problems common to all plant groups; the success in using DNA barcoding is lineage-dependent (Li et al. 2015). Identifying herbarium specimens using this method is also more difficult compared to fresh material, since dried specimens require a greater combination of primers that increases the chances of incorrect sequencing (Li et al. 2015; Vere et al. 2012). Furthermore, the widespread use of this technique is still limited because of the high cost (Stein et al. 2014).

Alternatively, one of the most promising tools currently used in botanical identification is Fourier-transform near-infrared spectroscopy (FT-NIR) (e.g., Lang et al. 2017; Rodríguez-Fernandez et al. 2011). The principle of the technique is to irradiate fractions of biological material (e.g., a dry leaf) in the infrared region. As a result, a set of absorbance values at different wavelengths (the spectra) is defined for the material (Workman and Weyer 2007). The spectra reflect molecular bonds, such as C-H, N-H, S-H or O-H, and are therefore related to biological molecules and the metabolome of the irradiated tissue (Stuart 2005).

Research using near-infrared spectroscopy to discriminate plant species is gaining more and more attention in plant taxonomy, especially for angiosperms (Durgante et al. 2013; Kim et al. 2004; Krajšek et al. 2008; Lang et al. 2017). The tool has been shown to be more practical and accurate than genetic or morphological methods (Castillo et al. 2008), is capable of consistently discriminating phylogenetic relationships of flowering plant species (Kim et al. 2004), and has been used in different works to aid in species circumscription and identification of several plant groups (Damasco et al. 2019; Durgante et al. 2013; Lang et al. 2017; Prata et al. 2018; Shen et al. 2020). However, the technique has not been tested to identify other groups of embryophytes, such as ferns and lycophytes, or bryophytes. (Guzmán et al. 2020).

Ferns are the second most diverse group of vascular plants, occur from tundras to tropical forests, and occupy niches from the ground to the canopy (Moran 2008). Due to the absence of flowers, fruits, and seeds, fern identification relies mainly on rhizome, frond, and sorus morphology (Schoute 1938). Sporophyte characters such as indument (trichomes and scales), leaf shape, and the structure and arrangement of sori are fundamental elements in the differentiation between species (Christenhusz and Chase 2014; Schoute 1938).

*Microgramma* (Polypodiaceae) comprises ca. 30 species, occurs in the Neotropics and tropical Africa (Almeida 2014), and is monophyletic according to the most recent circumscriptions (Salino et al. 2008; Almeida et al. in press). The genus exhibits wide morphological variation, especially in the leaves (e.g., it has both monomorphic and dimorphic species), leaf indument, and sorus arrangement (Almeida 2014). Additionally, intraspecific phenotypic variation and interspecific morphological overlap are found in closely related species, and there are species complexes, which may result in misidentifications in the genus (Almeida et al. in press). Using *Microgramma* as a model, our goal was to test the effectiveness of Fourier-transform near-infrared spectroscopy (FT-NIR) at discriminating and identifying closely related species in the fern lineage.

## METHODS

### Sampling

Dried leaves were selected from specimens at the BHCB, HSTM, and INPA herbaria (acronyms according to Thiers 2020 onwards: http://sweetgum.nybg.org/science/ih/). One hundred specimens belonging to thirteen species of *Microgramma* were analyzed (Appendix): *M. baldwinii* Brade, *M. crispata* (Fée) R.M.Tryon & A.F.Tryon, *M. dictyophylla* (Kunze ex Mett.) de la Sota, *M. geminata* (Schrad.) R.M.Tryon & A.F.Tryon, *M. lindbergii* (Mett. ex Kuhn) de la Sota, *M. lycopodioides* (L.) Copel., *M. megalophylla* (Desv.) de la Sota, *M. percussa* (Cav.) de la Sota, *M. persicariifolia* (Schrad.) C.Presl, *M. reptans* (Cav.) A.R.Sm., *M. squamulosa* (Kaulf.) de la Sota, *M. thurnii* (Baker) R.M.Tryon & Stolze, and *M. vacciniifolia* (Langsd. & Fisch.) Copel. All specimens had their identification confirmed by an expert (senior author). Only species with a minimum of five available specimens, with both fertile and sterile leaves, were selected. When possible, samples with fronds that were very damaged by insects or with signs of fungi or other epiphilic organisms were avoided. Sixteen spectral readings were obtained for each specimen (when possible), which included four readings, two on the adaxial surface and two on the abaxial surface, of four different leaves. No distinction between fertile and sterile leaves was made. The acquisition of the spectra lasted 30 seconds per reading and was taken using a Thermo Nicollet spectrophotometer, FT-NIR Antaris II Method Development System (MDS). The spectral readings consisted of 1,557 leaf absorbance values in the region of 4,000 to 10,000 cm-1 (1000 to 2500 nm). Each measurement produced by the equipment was an average of 16 readings with a wavelength resolution of 8 cm-1. The equipment was calibrated every 4 hours of use. A black body was placed over the frond to prevent light scattering.

### Analyses

All analyses were implemented in the statistical program R version 4.0.2 (R Core Team 2020). Three datasets using all FT-NIR spectrum wavelengths were tested to construct the spectral models: data of (i) adaxial+abaxial surfaces, (ii) adaxial surface only, and (iii) abaxial surface only. The datasets were explored using a principal component analysis (PCA). This technique allows the visualization of data of a smaller set of variables but still preserves the maximum information from the original variable set (Hongyu et al. 2016), thus allowing an exploratory analysis of the behavior of the spectra. The results of the PCA were represented in two-dimensional graphs using the first two main components with higher variation in the data.

To predict species based on spectral data, we used two supervised pattern recognition techniques: linear discriminant analysis (LDA) and partial least squares discriminant analysis (PLS-DA) (Berrueta and Héberger 2007). The LDA is a technique that discriminates and classifies objects based on previously defined groups (Sharma and Paliwal 2015), where the dependent variables corresponded to the species (categories) and the independent variables represent the absorbance values in the near-infrared. The PLS-DA, which also classifies the samples according to defined categories, is based on finding components that better explain the variations of the variables between classes, giving less weight to the noise and uncorrelated variations (Mevik and Cederkvist 2004). Both models were tested using the three different datasets.

Cross-validation techniques were used to assess model performance and species discrimination. The K-fold validation technique (Burman 1989) is where the set of calibration samples is divided into K subsets, with a subset taken out for validation and the remaining K-1 subsets used to build the model. Thus, at the end of K steps, the data is used in both test subsets and validation (Yadav et al. 2016). Here we use K = 10, described as the value that presents the best performance in the sampling, with the least bias in the error rate estimates (Kohavi 1995).

The leave-one-out (LOO) technique uses k-1 samples to generate the discriminant function and the sample not included in the model serves to validate it, obtaining the percentage of the model’s prediction (Kohavi 1995). Thus, we compared the predictions of individual identities for each species in each of the datasets.

## RESULTS

We found considerable variation in the near-infrared spectral data among the sampled species (Fig. 1). Among the three datasets tested, the adaxial+abaxial (i) dataset showed 97.8 % of the spectral variation, the adaxial (ii) dataset showed 97.6 % of the spectral variation, and the abaxial (iii) dataset was the most representative with a spectral variation of 98.1 % (Fig. 2). For the abaxial (iii) dataset, individuals belonging to the same species tended to group more cohesively and consequently less mixed compared to the remaining two datasets (adaxial+abaxial [i] and adaxial [ii]) (Fig. 2).

**Figure 1.**
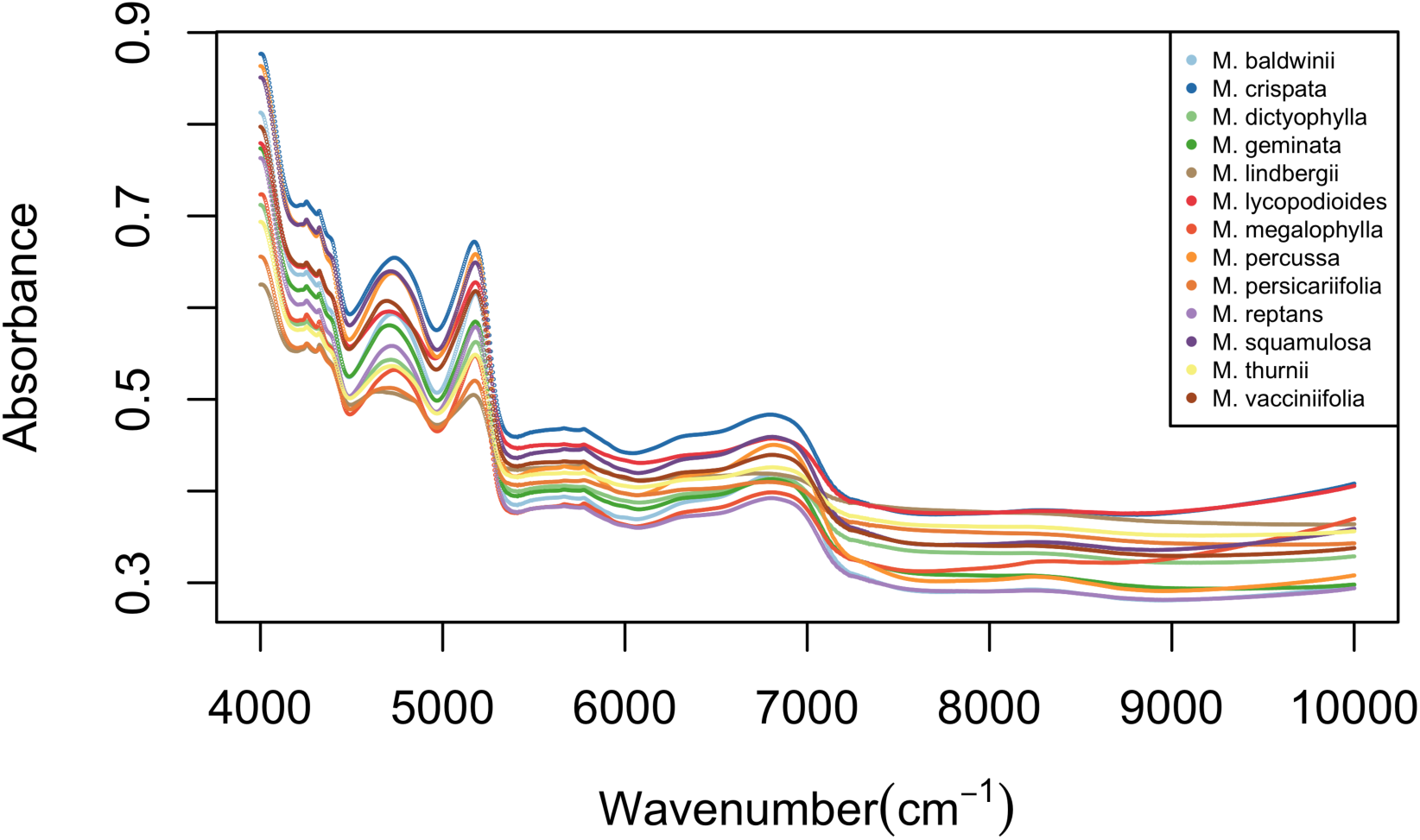
Average near-infrared spectral data for the thirteen sampled Microgramma species.

**Figure 2.**
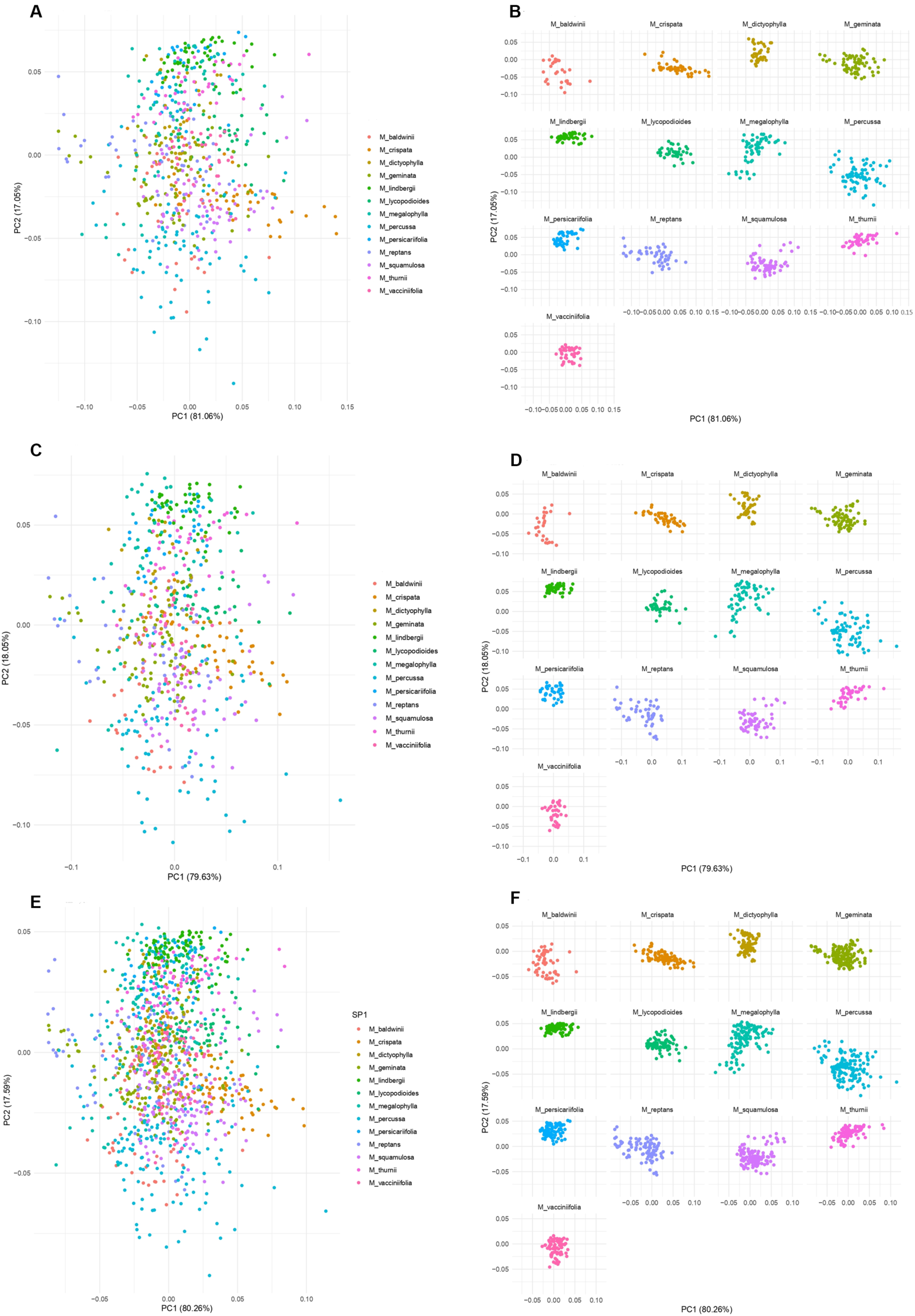
Principal component analysis (PCA) plot of the first two principal component axes for spectral data. **A.** Abaxial surface dataset (iii), all species. **B.** Abaxial surface dataset (iii), individually represented species. **C.** Adaxial surface dataset (ii), all species. **D.** Adaxial surface dataset (ii), individually represented species. **E.** Adaxial+abaxial surface dataset (i), all species. **F.** Adaxial+abaxial surface dataset (i), individually represented species.

All datasets had high predictive results in the identification of species (correct predictions higher than 90 %) for both models (PLS and LDA) and validation techniques (K-fold and LOO) (Table 1). Among the best percentages for plant discrimination (over 96 %) were the LDA model with the (iii) abaxial dataset for both the K-fold and leave-one-out validation techniques, and the PLS-DA model with the (i) adaxial+abaxial dataset and leave-one-out validation.

**Table 1.** Average percentage of correct identifications using a discriminant analysis and all of the FT-NIR spectrum wavelength data (1000–2500 nm) for the three datasets, (i) adaxial+abaxial leaf surfaces, (ii) adaxial surface only and (iii) abaxial surface only, for both models (LDA = linear discriminant analysis, PLS-DA = partial least squares discriminant analysis) and validation tests (K-fold and leave-one-out).

| Dataset | LDA |  | PLS-DA |  |
| --- | --- | --- | --- | --- |
|  | K-fold | LOO | K-fold | LOO |
| Adaxial+abaxial | 95.3 | 93.0 | 96.2 | 96.7 |
| Adaxial | 95.4 | 95.8 | 93.9 | 94.3 |
| Abaxial | 96.2 | 96.2 | 94.6 | 95.1 |

The adaxial+abaxial (i) dataset alone had the best percentage only for the PLS model and leave-one-out validation (96.7 %), and both validations in a similar way resulted in elevated correct identifications for the three datasets tested (adaxial+abaxial, adaxial, abaxial).

All individuals of *M. crispata* and *M. megalophylla* were 100 % correctly predicted in both models and validations tests, with no confusion of readings with any sample related to any other species (Figs 3, 4). For six species, *M. dictyophylla*, *M. geminata*, *M. lindbergii, M. lycopodioides*, *M. percussa*, and *M. reptans*, the correct prediction of the identities of individuals in all models and validations ranged from 90 to 100 % (Figs. 3, 4).

**Figure 3.**
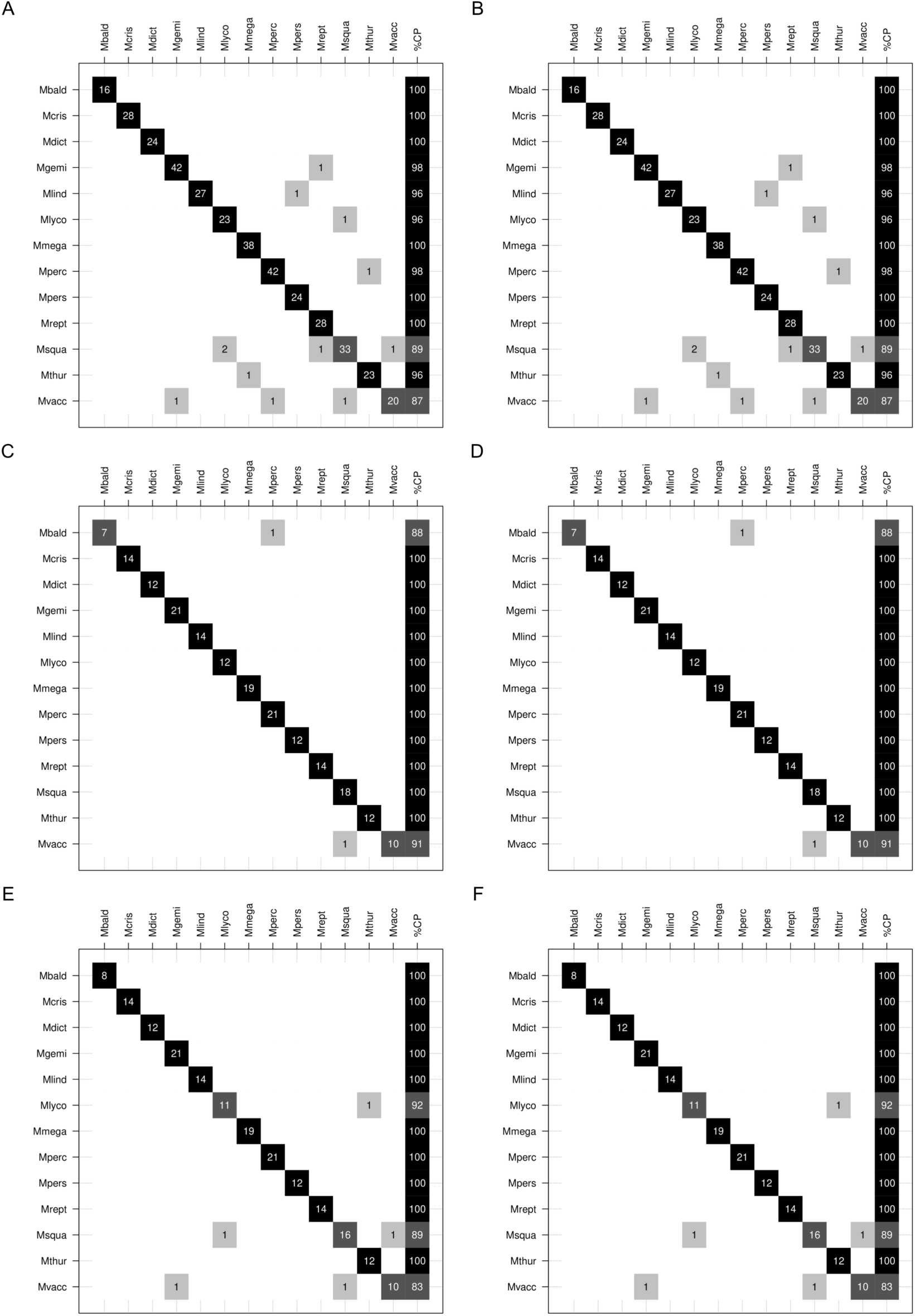
Confusion matrices resulting from the linear discriminant analysis (LDA) for the LOO and K-fold validations. (A) LOO validation, adaxial+abaxial surface data; (B) K-fold validation, adaxial+abaxial surface data; (C) LOO validation, abaxial surface data only; (D) K-fold validation, abaxial surface data only; (E) LOO validation, adaxial surface data only; (F) K-fold validation, adaxial surface data only. The names of the species observed are in rows and columns. The values on the diagonal correspond to correct predictions and those outside the diagonal correspond to incorrect predictions. Abbreviations: *M. bald* = M. baldwinii; *M. cris* = M. crispata; *M. dict* = M. dictyophylla; *M. gemi* = M. geminata; *M. lind* = M. lindbergii; *M. lyco* = M. lycopodioides; *M. mega* = M. megalophylla; *M. perc* = M. percussa; *M. pers* = M. persicariifolia; *M.rept* = M. reptans; *M. squa* = M. squamulosa; *M. thur* = M. thurnii; *M. vacc* = M. vacciniifolia.

**Figure 4.**
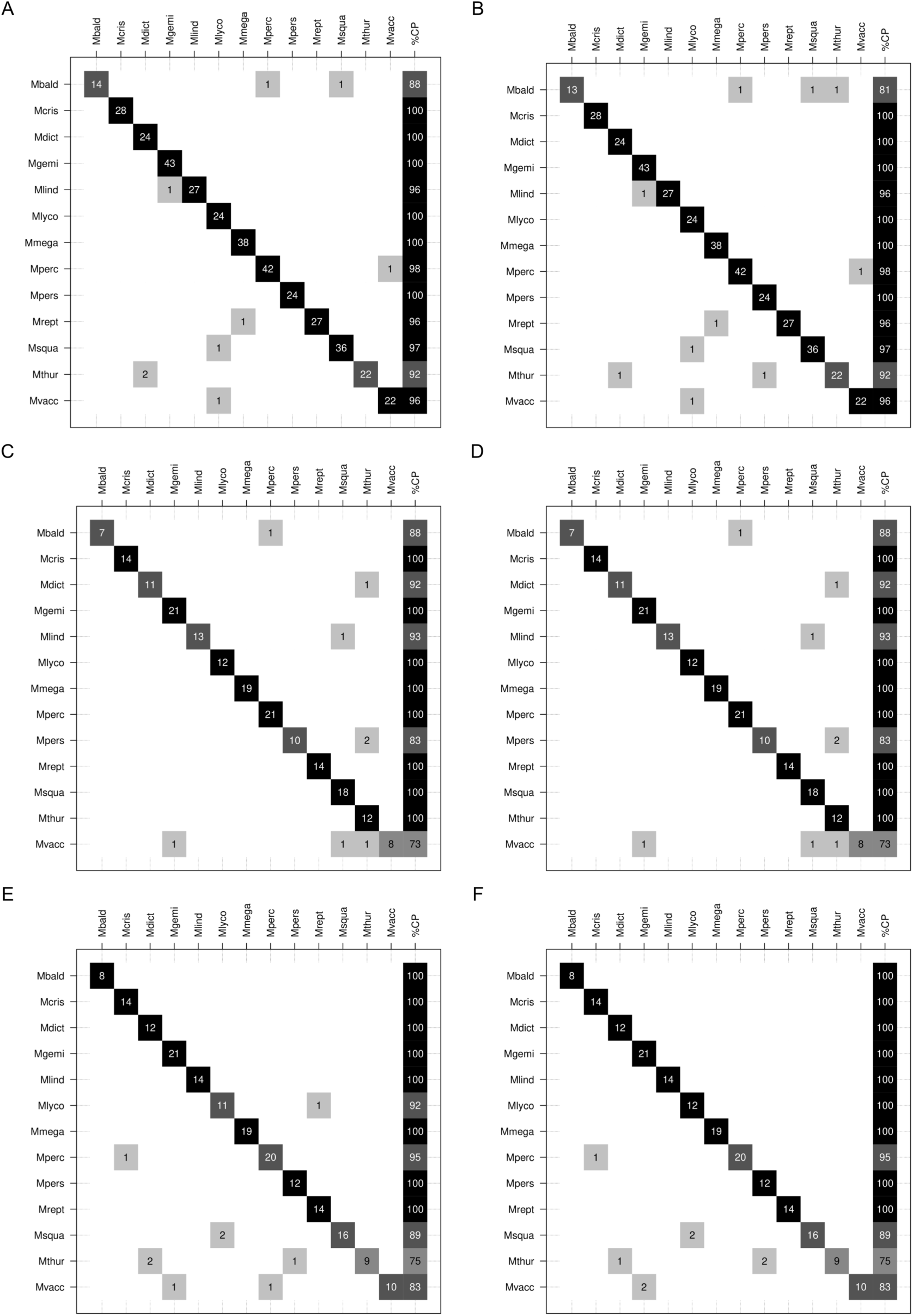
(A) K-fold validation, adaxial+abaxial surface data; (B) LOO validation, adaxial+abaxial surface data; (C) LOO validation, abaxial surface data only; (D) K-fold validation, abaxial surface data only; (E) K-fold validation, adaxial surface data only; (F) LOO validation, adaxial surface data only. The names of the species observed are in rows and columns. The values on the diagonal correspond to correct predictions and those outside the diagonal correspond to incorrect predictions. Abbreviations: *M. bald* = M. baldwinii; *M. cris* = M. crispata; *M. dict* = M. dictyophylla; *M. gemi* = M. geminata; *M. lind* = M. lindbergii; *M. lyco* = M. lycopodioides; *M. mega* = M. megalophylla; *M. perc* = M. percussa; *M. pers* = M. persicariifolia; *M.rept* = M. reptans; *M. squa* = M. squamulosa; *M. thur* = M. thurnii; *M. vacc* = M. vacciniifolia.

Two species (15.3 %), *M. persicariifolia* and *M. squamulosa*, had correct predictions between 80 and 100 % among the models and validations tests (Figs. 3, 4). For *M. baldwinii*, the abaxial (iii) dataset underperformed in both the LDA and PLS-DA models and validations (Figs. 3, 4). Additionally, in the PLS-DA model for this species, the adaxial+abaxial (i) dataset had 88 % and 81 % correct predictions for the K-fold and LOO validations, respectively. The remaining models and validations recovered 100 % correct predictions. For *M. persicariifolia*, the lowest prediction value (83 %) was found in the PLS model for the abaxial dataset, for both validations; the remaining models and validations recovered 100 % correct predictions. Regarding *M. squamulosa*, only the abaxial (iii) dataset had 100 % correct predictions in both models and validations (Figs. 3, 4).

The two species with the lowest percentages of correct predictions were *M. thurnii* and *M. vacciniifolia*. For *M. thurnii*, of the 12 different combinations of the datasets and tests, only two recovered one of the lowest percentages of correct predictions (75 %): the PLS model, with the adaxial (ii) dataset, for both validations. Four tests recovered more than 90 % correct predictions, and six had 100 % correct identifications (Figs. 3, 4). For this species, the PLS-DA models underperformed compared to the LDA models.

*Microgramma vacciniifolia* was the species with the lowest percentage of correct predictions (73 %), which was found by the PLS model with the adaxial+abaxial and adaxial datasets; although, for the abaxial dataset there was 100 % accuracy in the identifications (data not shown). Prediction errors for *M. vacciniifolia* individuals occurred mainly with spectra associated with *M. geminata* and *M. squamulosa* in both models. Even so, the lowest percentage observed was 73 % in the PLS model (Fig. 4).

## DISCUSSION

This is the first time that Fourier-transform near-infrared spectroscopy (FT-NIR) was tested for discriminating ferns species. Our results show that FT-NIR is a powerful tool that can be easily applied to species identification using spectral data of leaves. For all different scenarios tested (species, datasets, models, and validations), more than 85 % had an accuracy equal or greater than 90 % (Figs. 3, 4), with an average above 93 % (Table 1).

Regarding the accuracy of FT-NIR at species identification, we recognize the importance of using well-defined species circumscriptions and well-identified samples for constructing spectral models. In this work, when incorrectly identified specimens were used for the control group (an individual of *M. reptans* incorrectly determined as *M. baldwinii*), the accuracy decreased to 85.5 %, and after redoing the analysis with the correct identification, the correct prediction of the individuals of *M. baldwinii* reached 93.4 %.

The chemical composition and other structural characteristics of leaves vary within and between species, as a result of the developmental stage and a combination of environmental factors, ontogeny, and composition of the plant epidermis (Mediavilla et al. 2014). Ferns are characterized by the presence of sori, which are usually on the abaxial surface of the fronds (Schoute 1938). Some lineages exhibit leaf dimorphism, with leaves morphologically and physiologically specialized for photosynthesis or reproduction, and in some cases, there are extreme differences between both types (Wagner and Wagner 1977). In our study, we used species that are both monomorphic (*M. baldwinii, M. dictyophylla*, *M. geminata*, *M. lindbergii*, *M. lycopodioides*, *M. megalophylla*, *M. percussa*, *M. persicariifolia*, and *M. thurnii*) and dimorphic (*M. crispata*, *M. reptans*, *M. squamulosa*, and *M. vacciniifolia*). The dataset for the abaxial leaf surface, which can be more affected by the presence of sori, had (on average) higher percentages of discriminating samples than the other tested datasets (Table 1). Given our results, we believe the presence of sori has minimal influence on the spectral readings and subsequent discrimination power among species. However, this can vary among different lineages, and further tests controlling for fertile and sterile frond spectral readings are recommended.

One of the species that was more difficult to discriminate was *M. vacciniifolia*, where ca. 40 % of the samples were incorrectly predicted as *M. squamulosa*. These species are sympatric in eastern and central Brazil and exhibit wide morphological variation (Almeida 2014). Our results using near-infrared spectroscopy (NIR) could be revealing inconsistencies in their current taxonomic circumscriptions. Also, the existence of hybrids between these species (Sota 1973) might explain the related spectral readings and the lower percentage of correct predictions. The technique has been shown to detect differences in the physical and biochemical compositions expressed in plant samples, even between closely related species, populations, and hybrids (Atkinson et al. 1997; Cui et al. 2012; Humphreys et al. 2008).

Our best results show that the best models and validations can, on average, correctly predict the identification of species 96.7 % of the time when using all wavelengths to construct the models, which is comparable to previously published taxonomic works. Prata et al. (2018) demonstrated for the first time that near-infrared spectroscopy on leaves of subspecies of the *Pagamea guianensis* complex can discriminate taxa with high precision. Fan et al. (2010) tested and proved the reliability of the technique at discriminating *Ephedra* plants from different habitats and collection seasons, while Lang et al. (2017) showed the effectiveness of the technique at discriminating species, genera, and families of tree species from eighteen different angiosperm families. For groups of closely related plants, the technique has shown excellent results for species of *Protium* (Burseraceae), confirming the differences in spectral signatures among species (Damasco et al. 2019).

The high predictive power of FT-NIR at discriminating fern species, presented here, is superior to that observed for this lineage using a single region of two combined DNA barcodes, for which the best performance was 75 % correct predictions (Li et al. 2011; Wang et al. 2016). Identifying ferns and other plant groups using DNA barcoding is also an expensive technique and different lineages require specific combinations of molecular markers, which can make this technique complicated (Li et al. 2011; Lima et al. 2018). However, barcoding gametophytes has shown promising results for identifying species of ferns, which represents a great contribution to what is known about the evolution of this group (Schneider and Schuettpelz 2006). Our work does not intend to minimize the importance of other techniques used in plant systematics, but rather tested the reliability and effectiveness of FT-NIR at discriminating species in a group known to be problematic. Further, it highlights the potential of using this method in studies about plant systematics.

## CONCLUSION

Our results show that near-infrared spectroscopy (NIR) is a highly effective, cost-effective, and non-destructive technique that can be used to discriminate closely related species. In addition to the possibility of obtaining spectral data quickly with minimal damage to samples, the technique provides greater reliability at discriminating morphologically similar fern species, as previously found for some angiosperms. The accuracy of the identifications is comparable to and even surpasses that of DNA barcoding, even for species from highly diverse and heterogeneous areas, such as tropical forests. We believe that NIR has great potential to be used in integrative taxonomic studies that aim to better understand species circumscriptions in the fern lineage.

## ACKNOWLEDGMENTS

This study was partly financed by the Coordenação de Aperfeiçoamento de Pessoal de Nível Superior - Brasil (CAPES) – financing code 001, and Programa Nacional de Cooperação Acadêmica na Amazônia (PROCAD-AM/CAPES 21/2018, nº 88887.200472/2018-00). The authors thank the following: B. Leal, L.L. Giacomin, M.A. Buitrago, and T. André for contributing to the manuscript; Mike Hopkins for all the support given to the first author while working at INPA; the herbaria that kindly gave us access to the specimens used in this research; and our colleagues from HSTM and INPA for the support and willingness to help.

## APPENDIX

Specimens used for spectral data capture. Herbaria acronym (in parentheses) follow Thiers (2020 onwards: http://sweetgum.nybg.org/science/ih/).

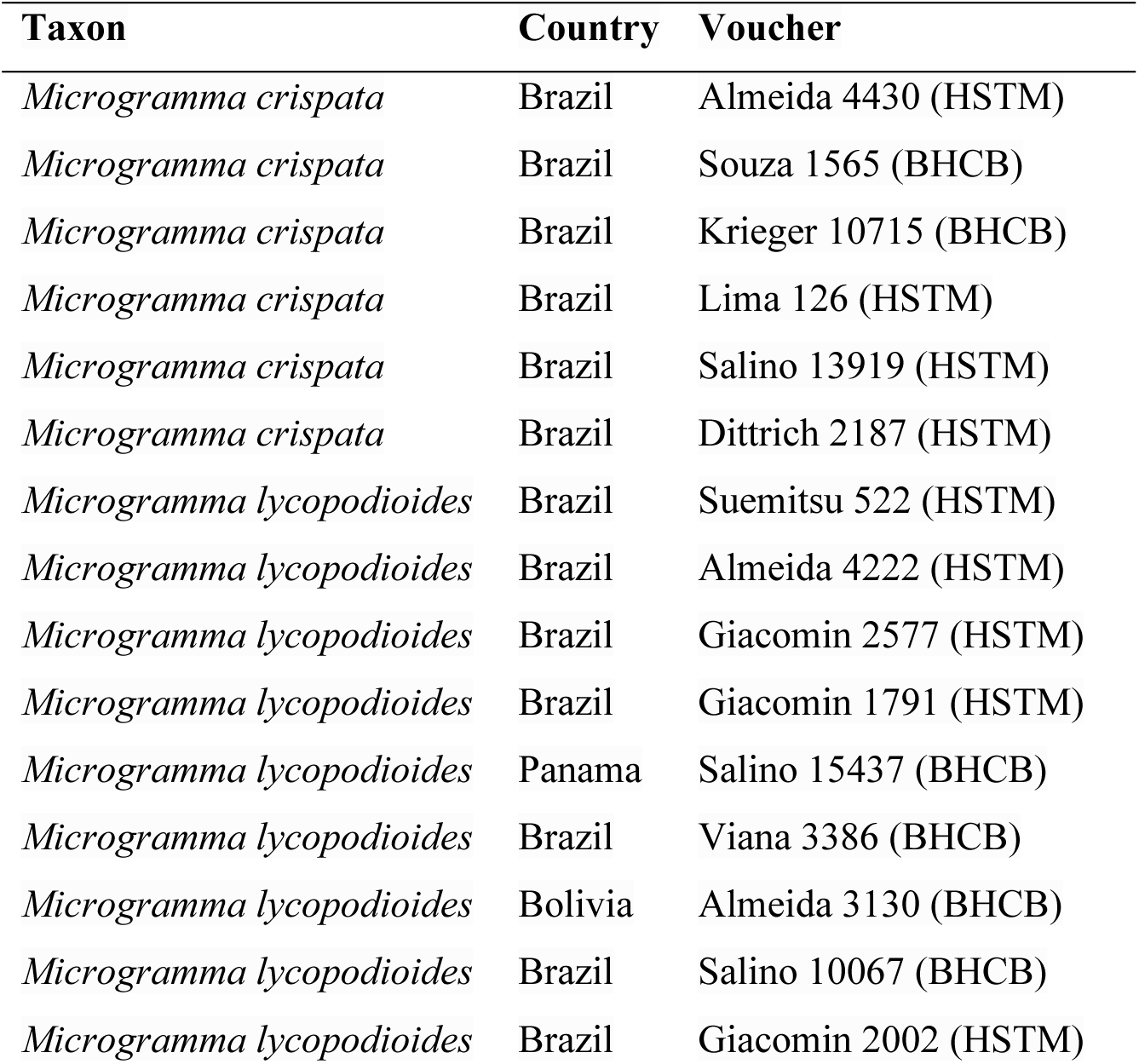
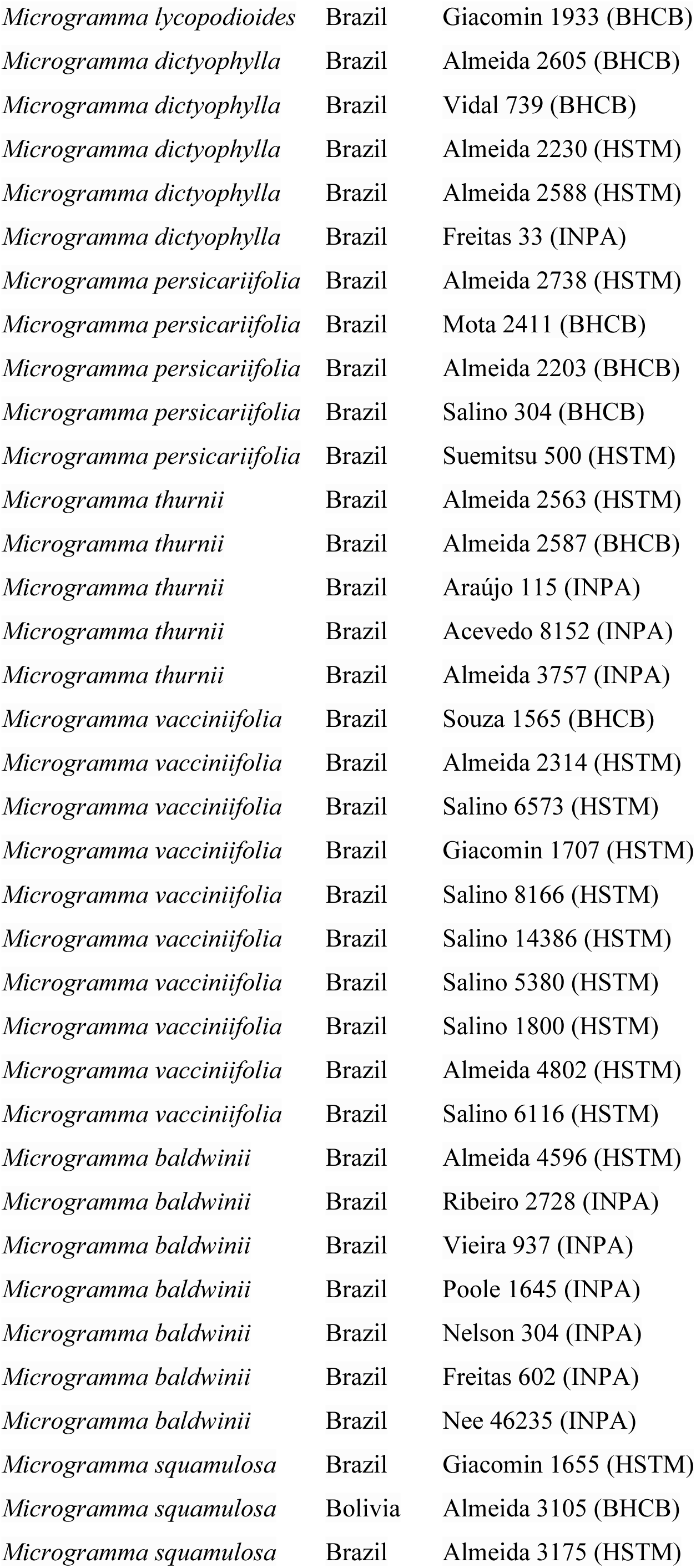
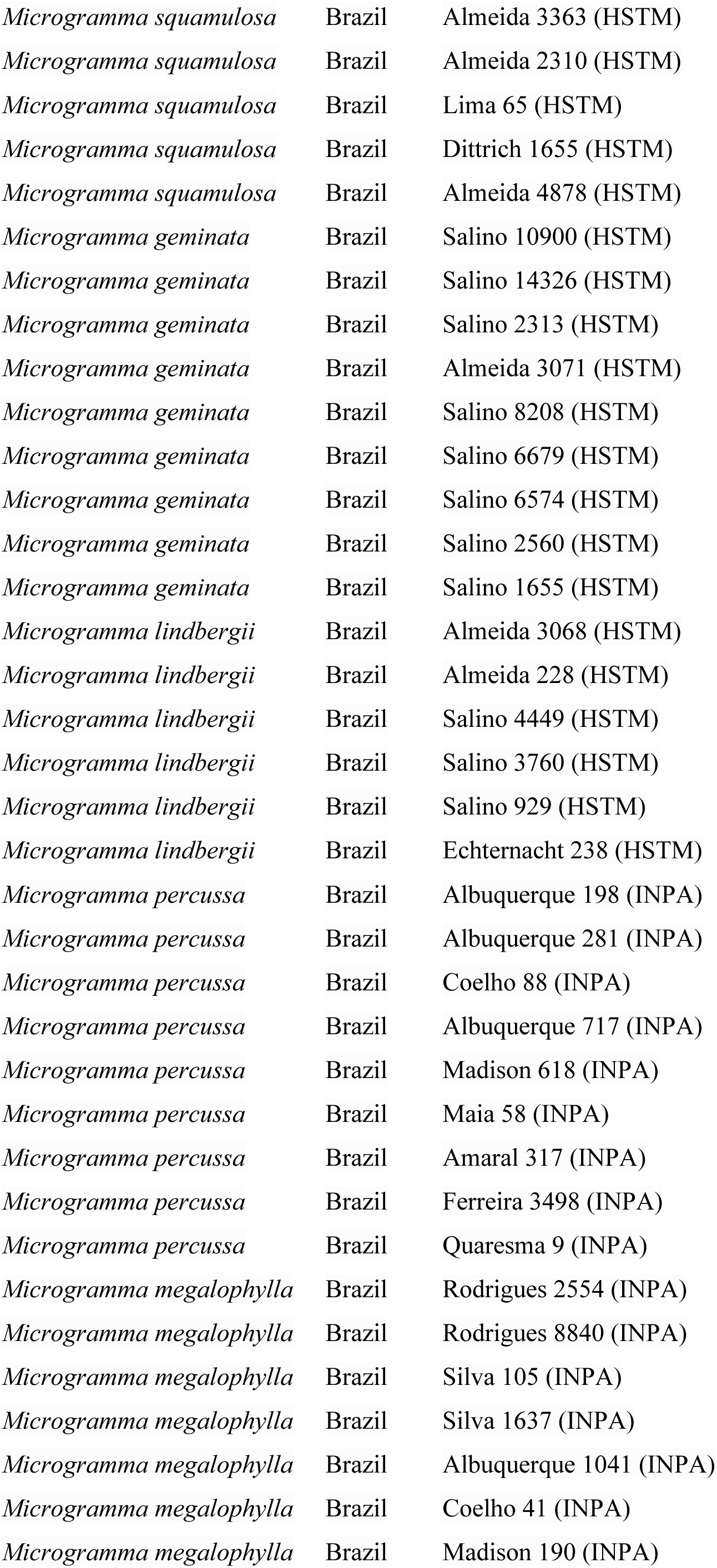
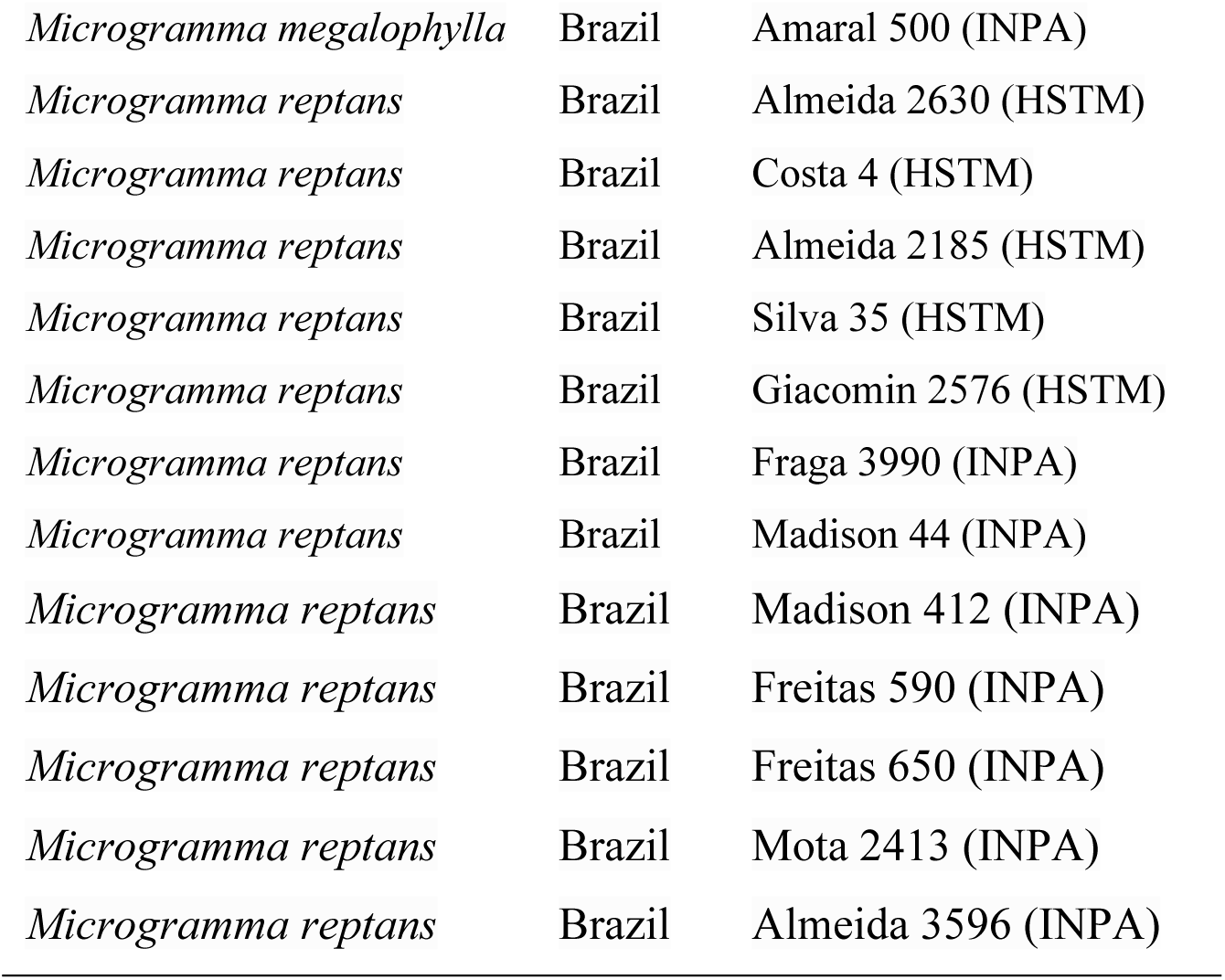

| Taxon | Country | Voucher |
| --- | --- | --- |
| *Microgramma crispata* | Brazil | Almeida 4430 (HSTM) |
| *Microgramma crispata* | Brazil | Souza 1565 (BHCB) |
| *Microgramma crispata* | Brazil | Krieger 10715 (BHCB) |
| *Microgramma crispata* | Brazil | Lima 126 (HSTM) |
| *Microgramma crispata* | Brazil | Salino 13919 (HSTM) |
| *Microgramma crispata* | Brazil | Dittrich 2187 (HSTM) |
| *Microgramma lycopodioides* | Brazil | Suemitsu 522 (HSTM) |
| *Microgramma lycopodioides* | Brazil | Almeida 4222 (HSTM) |
| *Microgramma lycopodioides* | Brazil | Giacomin 2577 (HSTM) |
| *Microgramma lycopodioides* | Brazil | Giacomin 1791 (HSTM) |
| *Microgramma lycopodioides* | Panama | Salino 15437 (BHCB) |
| *Microgramma lycopodioides* | Brazil | Viana 3386 (BHCB) |
| *Microgramma lycopodioides* | Bolivia | Almeida 3130 (BHCB) |
| *Microgramma lycopodioides* | Brazil | Salino 10067 (BHCB) |
| *Microgramma lycopodioides* | Brazil | Giacomin 2002 (HSTM) |
| *Microgramma lycopodioides* | Brazil | Giacomin 1933 (BHCB) |
| *Microgramma dictyophylla* | Brazil | Almeida 2605 (BHCB) |
| *Microgramma dictyophylla* | Brazil | Vidal 739 (BHCB) |
| *Microgramma dictyophylla* | Brazil | Almeida 2230 (HSTM) |
| *Microgramma dictyophylla* | Brazil | Almeida 2588 (HSTM) |
| *Microgramma dictyophylla* | Brazil | Freitas 33 (INPA) |
| *Microgramma persicariifolia* | Brazil | Almeida 2738 (HSTM) |
| *Microgramma persicariifolia* | Brazil | Mota 2411 (BHCB) |
| *Microgramma persicariifolia* | Brazil | Almeida 2203 (BHCB) |
| *Microgramma persicariifolia* | Brazil | Salino 304 (BHCB) |
| *Microgramma persicariifolia* | Brazil | Suemitsu 500 (HSTM) |
| *Microgramma thurnii* | Brazil | Almeida 2563 (HSTM) |
| *Microgramma thurnii* | Brazil | Almeida 2587 (BHCB) |
| *Microgramma thurnii* | Brazil | Araújo 115 (INPA) |
| *Microgramma thurnii* | Brazil | Acevedo 8152 (INPA) |
| *Microgramma thurnii* | Brazil | Almeida 3757 (INPA) |
| *Microgramma vacciniifolia* | Brazil | Souza 1565 (BHCB) |
| *Microgramma vacciniifolia* | Brazil | Almeida 2314 (HSTM) |
| *Microgramma vacciniifolia* | Brazil | Salino 6573 (HSTM) |
| *Microgramma vacciniifolia* | Brazil | Giacomin 1707 (HSTM) |
| *Microgramma vacciniifolia* | Brazil | Salino 8166 (HSTM) |
| *Microgramma vacciniifolia* | Brazil | Salino 14386 (HSTM) |
| *Microgramma vacciniifolia* | Brazil | Salino 5380 (HSTM) |
| *Microgramma vacciniifolia* | Brazil | Salino 1800 (HSTM) |
| *Microgramma vacciniifolia* | Brazil | Almeida 4802 (HSTM) |
| *Microgramma vacciniifolia* | Brazil | Salino 6116 (HSTM) |
| *Microgramma baldwinii* | Brazil | Almeida 4596 (HSTM) |
| *Microgramma baldwinii* | Brazil | Ribeiro 2728 (INPA) |
| *Microgramma baldwinii* | Brazil | Vieira 937 (INPA) |
| *Microgramma baldwinii* | Brazil | Poole 1645 (INPA) |
| *Microgramma baldwinii* | Brazil | Nelson 304 (INPA) |
| *Microgramma baldwinii* | Brazil | Freitas 602 (INPA) |
| *Microgramma baldwinii* | Brazil | Nee 46235 (INPA) |
| *Microgramma squamulosa* | Brazil | Giacomin 1655 (HSTM) |
| *Microgramma squamulosa* | Bolivia | Almeida 3105 (BHCB) |
| *Microgramma squamulosa* | Brazil | Almeida 3175 (HSTM) |
| *Microgramma squamulosa* | Brazil | Almeida 3363 (HSTM) |
| *Microgramma squamulosa* | Brazil | Almeida 2310 (HSTM) |
| *Microgramma squamulosa* | Brazil | Lima 65 (HSTM) |
| *Microgramma squamulosa* | Brazil | Dittrich 1655 (HSTM) |
| *Microgramma squamulosa* | Brazil | Almeida 4878 (HSTM) |
| *Microgramma geminata* | Brazil | Salino 10900 (HSTM) |
| *Microgramma geminata* | Brazil | Salino 14326 (HSTM) |
| *Microgramma geminata* | Brazil | Salino 2313 (HSTM) |
| *Microgramma geminata* | Brazil | Almeida 3071 (HSTM) |
| *Microgramma geminata* | Brazil | Salino 8208 (HSTM) |
| *Microgramma geminata* | Brazil | Salino 6679 (HSTM) |
| *Microgramma geminata* | Brazil | Salino 6574 (HSTM) |
| *Microgramma geminata* | Brazil | Salino 2560 (HSTM) |
| *Microgramma geminata* | Brazil | Salino 1655 (HSTM) |
| *Microgramma lindbergii* | Brazil | Almeida 3068 (HSTM) |
| *Microgramma lindbergii* | Brazil | Almeida 228 (HSTM) |
| *Microgramma lindbergii* | Brazil | Salino 4449 (HSTM) |
| *Microgramma lindbergii* | Brazil | Salino 3760 (HSTM) |
| *Microgramma lindbergii* | Brazil | Salino 929 (HSTM) |
| *Microgramma lindbergii* | Brazil | Echternacht 238 (HSTM) |
| *Microgramma percussa* | Brazil | Albuquerque 198 (INPA) |
| *Microgramma percussa* | Brazil | Albuquerque 281 (INPA) |
| *Microgramma percussa* | Brazil | Coelho 88 (INPA) |
| *Microgramma percussa* | Brazil | Albuquerque 717 (INPA) |
| *Microgramma percussa* | Brazil | Madison 618 (INPA) |
| *Microgramma percussa* | Brazil | Maia 58 (INPA) |
| *Microgramma percussa* | Brazil | Amaral 317 (INPA) |
| *Microgramma percussa* | Brazil | Ferreira 3498 (INPA) |
| *Microgramma percussa* | Brazil | Quaresma 9 (INPA) |
| *Microgramma megalophylla* | Brazil | Rodrigues 2554 (INPA) |
| *Microgramma megalophylla* | Brazil | Rodrigues 8840 (INPA) |
| *Microgramma megalophylla* | Brazil | Silva 105 (INPA) |
| *Microgramma megalophylla* | Brazil | Silva 1637 (INPA) |
| *Microgramma megalophylla* | Brazil | Albuquerque 1041 (INPA) |
| *Microgramma megalophylla* | Brazil | Coelho 41 (INPA) |
| *Microgramma megalophylla* | Brazil | Madison 190 (INPA) |
| *Microgramma megalophylla* | Brazil | Amaral 500 (INPA) |
| *Microgramma reptans* | Brazil | Almeida 2630 (HSTM) |
| *Microgramma reptans* | Brazil | Costa 4 (HSTM) |
| *Microgramma reptans* | Brazil | Almeida 2185 (HSTM) |
| *Microgramma reptans* | Brazil | Silva 35 (HSTM) |
| *Microgramma reptans* | Brazil | Giacomin 2576 (HSTM) |
| *Microgramma reptans* | Brazil | Fraga 3990 (INPA) |
| *Microgramma reptans* | Brazil | Madison 44 (INPA) |
| *Microgramma reptans* | Brazil | Madison 412 (INPA) |
| *Microgramma reptans* | Brazil | Freitas 590 (INPA) |
| *Microgramma reptans* | Brazil | Freitas 650 (INPA) |
| *Microgramma reptans* | Brazil | Mota 2413 (INPA) |
| *Microgramma reptans* | Brazil | Almeida 3596 (INPA) |

## Notes

### Competing Interest Statement

The authors have declared no competing interest.

